# Fine temporal brain network structure modularizes and localizes differently in men and women: Insights from a novel explainability framework

**DOI:** 10.1101/2022.06.09.495551

**Authors:** Noah Lewis, Robyn Miller, Harshvardhan Gazula, Vince Calhoun

## Abstract

Deep learning has become an effective tool for classifying biological sex based on functional magnetic resonance imaging (fMRI), but research on what features within the brain are most relevant to this classification is still lacking. Model interpretability has become a powerful way to understand “black box” deep-learning models and select features within the input data that are most relevant to the correct classification. However, very little work has been done employing these methods to understand the relationship between the temporal dimension of functional imaging signals and classification of biological sex, nor has there been attention paid to rectifying problems and limitations associated with feature explanation models, e.g. underspecification and instability. We provide a methodology to limit the impact of underspecification on the stability of the measured feature importance, and then, using intrinsic connectivity networks (ICNs) from fMRI data, we provide a deep exploration of sex differences among functional brain networks. We report numerous conclusions, including activity differences in the visual and cognitive domains, as well as major connectivity differences.

## 1 Introduction

Deep learning has been shown to be an effective tool for both classification of biological sex as well as understanding the features relevant to the classification [1, 2, 3, 4]. However, it suffers from two critical flaws from the standpoint of model interpretability: underspecification and instability of the relevant features. Underspecified [5] models can have many local minima, or possible functions, which produce the same mapping between the input and the output under different parameters. This is particularly problematic for feature attribution methods such as saliency [6], which are very sensitive to changes in model architecture, even to initialization within a given architecture. Although saliency methods can be informative about the data, there are several technical issues. Primarily, they are often unstable and very sensitive to small perturbations of the model architecture or initialization state. This instability is particularly pronounced for deep classification models applied to small datasets. A secondary flaw of saliency is specific to sequential/recurrent models, such as long-short term memory models (LSTMs). In this case, saliency methods become ineffective due to a phenomenon known as vanishing saliency [7], which significantly reduces the magnitude of the salient gradients as the model backpropagates through time, providing inaccurate saliency maps.

In recent decades, functional magnetic resonance imaging (fMRI), has significantly extended our understanding of the human brain [8]. We have witnessed great strides in analyzing fMRI data, particularly through the use of independent component analysis (ICA) to extract intrinsic connectivity networks (ICNs) [9]. These ICNs and their associated timecourses have become central to fMRI research, including research into brain activation patterns and biological sex. Although there is now a wealth of information about sex differences among brain signals, there is still a long way to go before we truly understand the nuances of how brain signals relate to biological sex [10]. One of the more promising avenues for fMRI research is the analysis of complex brain disorders such as schizophrenia, Alzheimer’s, and autism. As a great deal of research has found sex differences relating to these disorders [11, 12, 13], it is imperative to better understand the relationship between sex and fMRI signals as a whole. With a better understanding of this relationship, researchers may have a better grasp of how best to treat these disorders based on sex. This can include medications, dosage, and behavioral treatments.

In this paper, we present a methodology to mitigate instability in feature importance assessments using state-of-the-art, non-linear models and feature attribution methods, then apply these methodologies to elucidate the relationship between biological sex and mesoscale brain dynamics. Specifically, using an LSTM model coupled with a specific saliency method known as integrated gradients (IG), we take a deep-dive into understanding sex differences among functional networks estimated from fMRI data. LSTMs are important for this work because they can capture the dynamics of fMRI temporal signals. Lastly, we present evidence that deep learning can be usefully employed as a “feature explainer”, or a tool which highlights aspects of the brain function most relevant to sex differences.

With the power of novel deep learning methods, we take an in-depth look at sex differences among fMRI ICN timecourses. These timecourses represent interpretable functional networks, making quantitative analysis of our results relatively easy. We also use a very large sample size, from the UK Biobank (UKB) repository, which aids both interpretability and model stability. Our approach to investigating these networks is a novel adaptation of simple feature explanation techniques that fixes several key problems, primarily the instability of the feature maps and an LSTM-specific issue, a phenomenon known as vanishing saliency. We then validate our methodology with synthetic data in which the most relevant features are known beforehand. While showing visual representations of our maps, we also quantitatively compare our proposed methodology to an existing methodology, the input-cell attention [7]. We performed these analyses to ground our work and show quantitatively that our methods can discover truly data-relevant signals. After this validation, we provide a broad set of post-hoc analysis, showing both validity of our model as well as novel results, further expanding biological sex analysis based on fMRI data. Pointedly, we find, after comparing to static functional network connectivity (sFNC), considerable sex-specific results within relationships between the individual ICNs. And in particular, differences within key functional domains including the visual (VIS) and default mode network (DMN).

## 2 Data & Methods

### 2.1 FMRI data and preprocessing

The data, a total of 8216 (4202 males and 4014 females) resting-state fMRI scans, was sourced from across 22 sites within the United Kingdom between 2006 and 2018. Data processing and quality control were previously performed in [14], where over 11k subjects were selected from the entire UKB dataset. However, we removed subjects with inconclusive sex, meaning any subject where the documented biological sex is not consistent with the documented gender, or subjects missing either field, which gives us the final 8216 subjects. Participants were between 40 and 69 years of age [15, 16]. The data acquisition protocol is as follows: 39ms echo time, a 0.735 second repetition time (TR), 52^°^ flip angle, and a multiband factor of 8 using 3T Siemens machines. The T2 signal was both linearly and non-linearly warped to MNI152 space. Each volume was resampled to 3mm^3^ for a final image size of 53 × 63 × 46 mm^3^, and 160 time steps using the statistical parametric mapping (SPM12, http://www.fil.ion.ucl.ac.uk/spm/) MATLAB package.

After the preprocessing pipeline, the intrinsic connectivity networks (ICNs) were extracted using spatially constrained ICA with the GIFT package [17] via the Neuromark pipeline [18] for MATLAB. ICA is a robust and evidence-based method to capture regions of functional activity [19]. Since our goal is to analyze the brain functionality, we require clean and data-driven representations of this activity. We also want to compare the functional relationships measured with our methods to other connectivity metrics. Logically, we need the network timecourses so we can compare our analyses of the relationships between networks to other robust connectivity estimations. This pipeline provides a fully automated approach to compute ICA (both spatial components and timecourses) and output labeled and ordered components. Overall, there were 53 networks, covering 7 domains: Subcortical (SC), auditory (AUD), sensorimotor (SM), visual (VIS), cognitive control (CC), the default mode network (DMN), and the cerebellum (CB).

### 2.2 Our Model

A key aspect of our model, that mitigates vanishing saliency, is an additive attention mechanism [20] between the outputs of a bi-directional LSTM (bi-LSTM) [21] and the final output layer, which creates a direct gradient flow path from the classification to the input via the attention parameters [22]. A diagram of our model can be seen in figure 1. A bi-directional LSTM was chosen because we do not consider streaming data, the additional parameters aid training, and the extra directional flow for gradients may also improve the quality of the saliency maps.

**Figure 1:**
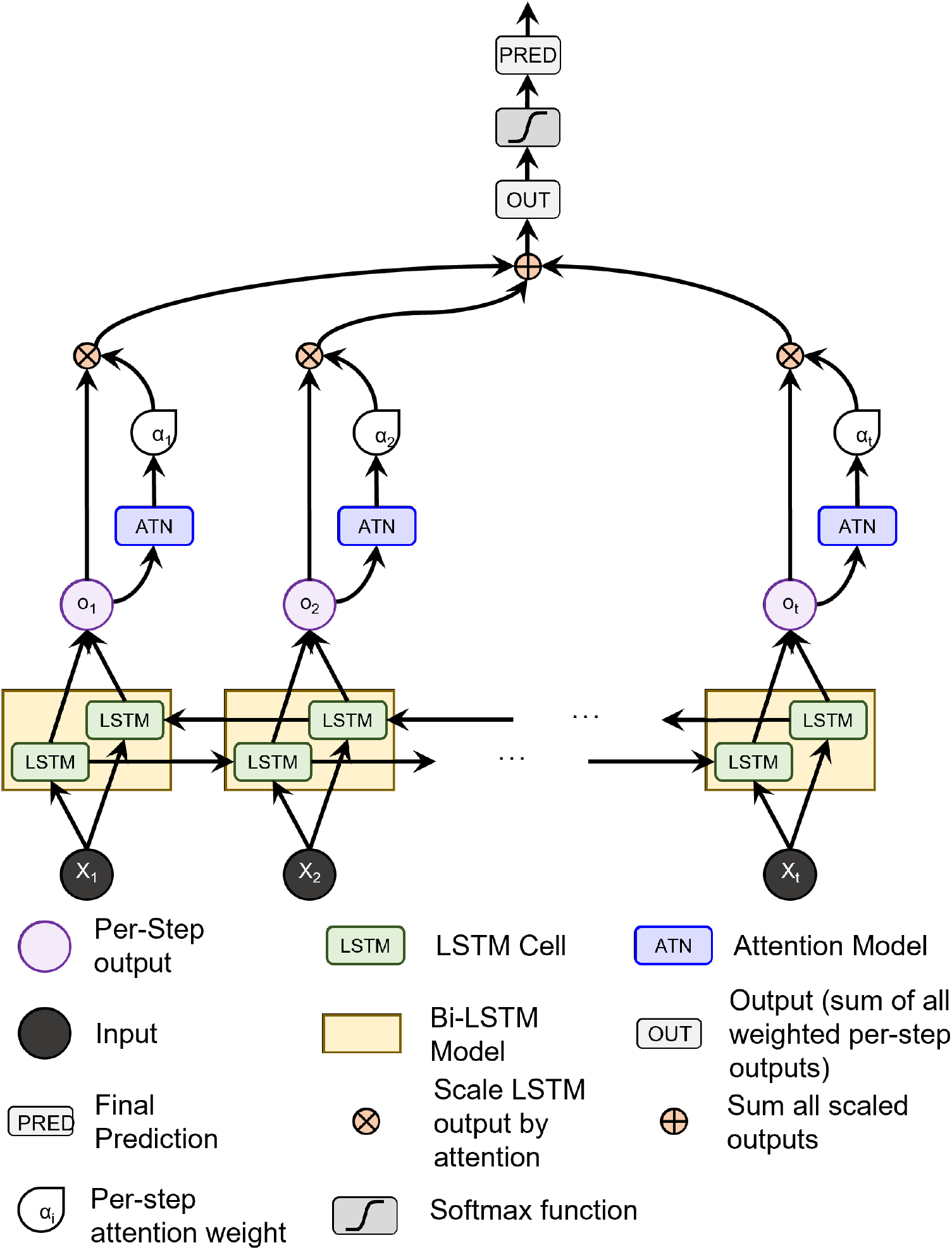
A diagram of our model. The data passes through an LSTM, from which the hidden states parameterize the attention model. The attention weights scale the hidden states, which are summed over all states (i.e. step in the LSTM), and is finally fed through a classification layer.

The attention mechanism [20] is a powerful way to amalgamate temporal information and “attend” only to the most important steps in the LSTM output by assigning a weight to each step. To parameterize the attention mechanism, we pass the LSTM output at each step through an attention network of two feed-forward layers to create a single, per-step weight value. The weight values from all time steps are jointly softmaxed and used to adjust the LSTM output at the individual time steps. As the model is bi-directional, we use the output from both the forward and backward directions concatenated into a single vector as our context for the attention mechanism. In other words, *h_backward*_*T*_ is concatenated with *h_forward*_*T*_ and passed through the attention mechanism to give us the respective attention weight for that time step. Once the hidden outputs have been individually weighted by the attention scalar, they are summed along the time dimension and pushed through a linear transform for classification.

### 2.3 Gradient-based Feature Attribution

Gradient-based feature attribution methods, commonly known as saliency, are model interpretability methods that leverage the gradients of a trained model to better understand why the model makes its predictions. It is defined as the gradients of the prediction of the correct class w.r.t. the input, or 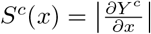.

With this paradigm of using calculated gradients to interpret and understand the input data, there are numerous methods to compute the gradients as 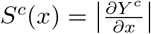. For our purposes, we chose a method called integrated gradients (IG) [23]. IG is defined, for any model, *F* as 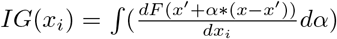, where *x*_*i*_ is feature *i* for a given input sample, *x*, and *x*^′^ is defined as a baseline image, where, for our purposes we use a zero-valued “blank” image as our baseline. Associated with the baseline image is an interpolation constant, *α*. Essentially, for each feature, we integrate over our interpolation factor, *α* over a function in which we subtract the baseline from the input, pass this modified input through the model, F, and then compute the gradients of the correct class with respect to input feature *i*. The interpolation factor is, in our case, a set of linear steps between the baseline image and the input. An example of these maps can be seen in figure 2. One thing to note about our implementation of integrated gradients is that we do not element-wise multiply our gradient maps with the input, which is unlike the original formulation. We do this to ensure no information from the input is enforced upon the gradient maps, meaning the attribution assignments from our maps are totally separate from the input itself. All of our feature attribution calculations come from the Captum python library [24].

**Figure 2:**
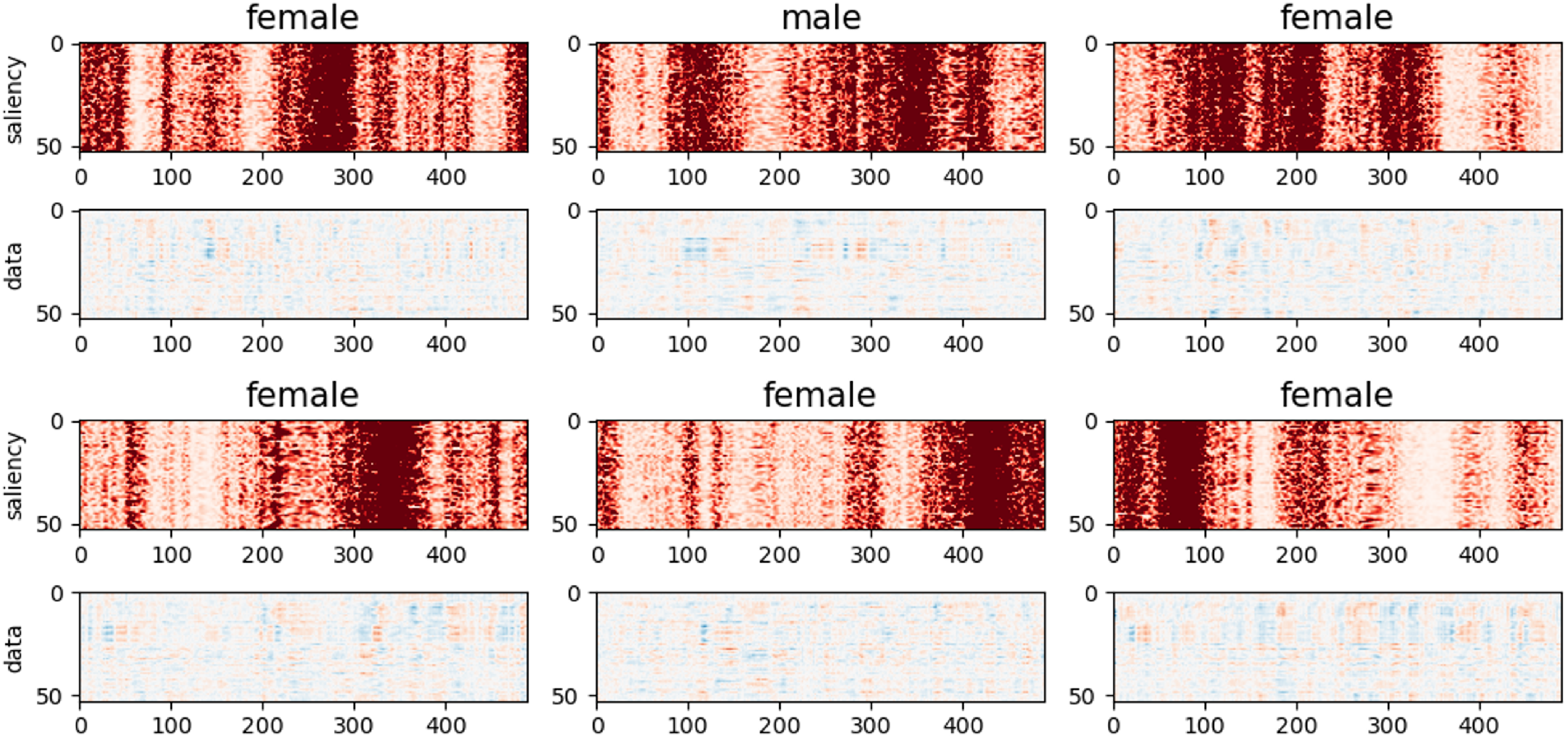
Saliency maps (red rows) from several subjects compared to their associated ICA time course (below).

### 2.4 Our Approach

Using 10-fold cross-validation, from the trained model of each fold, we calculate the IG maps (aka saliency maps) for each test sample. This accumulates to one map for each subject when the subject was used as a test sample. This methodology ensures that none of the maps were from training subjects, which could bias the resulting maps. These saliency maps highlight the features that the prediction likelihood of the correct class is the most sensitive to. However, we observe that the maps are rarely stable, and vary widely with the initial randomization. To correct for this, we train multiple models (keeping the hyperparameter settings the same) with different random initializations and calculate saliency maps from each model (for all experiments, we train 300 total models). For each input sample, we select the map that is closest (the distances are weighted by the loss for that subject/model pair to give more importance to better performing models) to the average map over all models for that sample, using Euclidean distance. Then, the selected maps are normalized by the sum. A diagram of our methodological flow is in figure 3.

**Figure 3:**
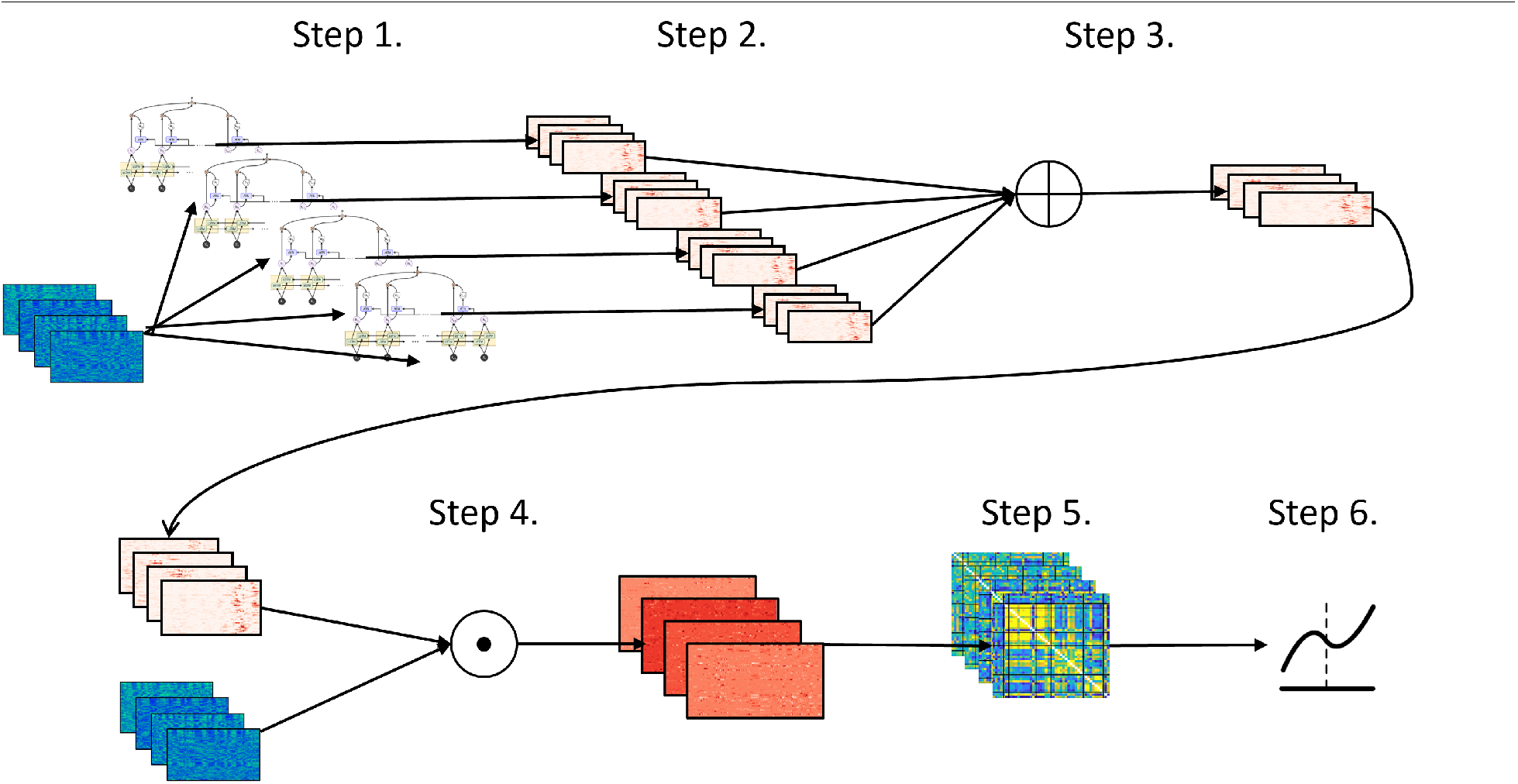
Flowchart describing our pipeline for analyzing the ICA timecourses. For all other datasets, we use only the first 4 steps to calculate the finalized maps. Step 1: we train 300 separate models (each with the same architecture) using different random initializations for each model on the same set of ICA timecourses. Step 2: we calculate the saliency maps for each sample from all 300 models. Step 3: We calculate the average saliency map for each sample over all 300 models. Step 4: We select the per-model set of maps with the lowest Euclidean distance to the average over all models, resulting in a stable saliency map for each input sample. This distance is weighted by the loss of that subject/model pair, giving more importance (shorter distances) to more accurate model/subject pairs.

### 2.5 Synthetic Data

The synthetic data is specifically engineered so that the relevant information within the data is quantifiable and interpretable. In this work, we use two sets of synthetic data, and a third experiment can be found in the supplementary material. In the first dataset of 30,000 samples, each sample is generated as random Gaussian noise with a sequence length of 200 and 30 features, then randomly assigned a class label of either 0 or 1. For each sample, a window of 10 time steps is randomly chosen, within which a portion of the data is perturbed based on the assigned label. If the label is 0, the first 15 features in each of the 10 time steps are perturbed, and if the label is 1, the last 15 features are perturbed. Each target feature is perturbed by adding randomly generated Gaussian noise. This creates a pattern of “boxes” for the dataset, an example of which can be seen in figure 4. In essence, only the features themselves are predictive of the class label, as opposed to temporal patterns. This box dataset, a trivial example, is a way to show the effectiveness of our methodology in a vacuum with very few confounding variables.

**Figure 4:**
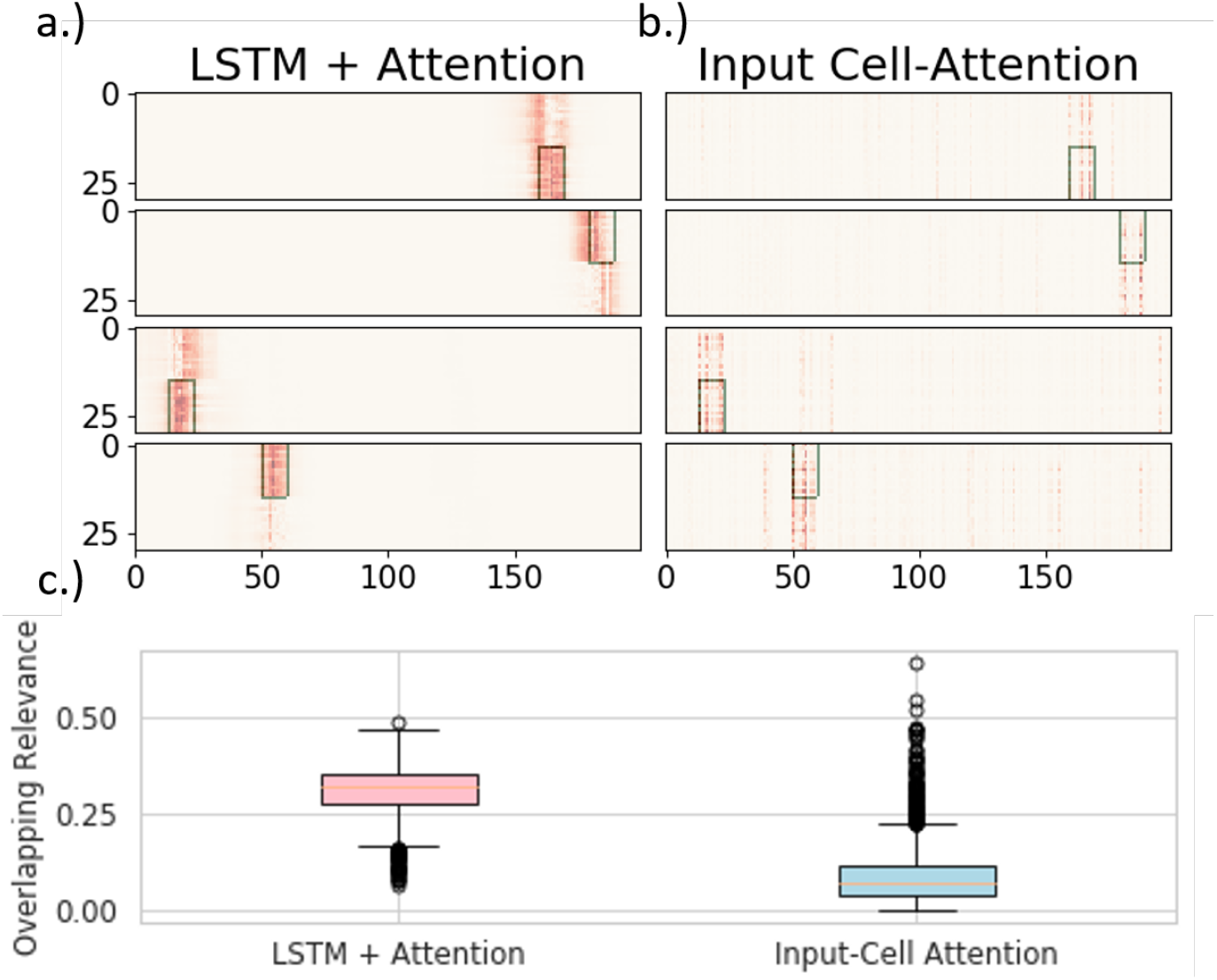
The results from the analysis of the boxes dataset. Several examples of resulting maps from both LSTM+attention (a) and input-cell attention (b) where the green rectangle is a mask representing the truly rele-vant information (i.e. box location). Boxplots (c) of the overlapping-values metric over all 3,000 samples for both models. The overlap is defined as the percentage of the total sum of the maps that are within the relevant area seen in the top figures.

A second synthetic experiment is used to show that the saliency maps are still accurate when the relevant information is based on dynamic patterns. As with the first experiment, each sample begins as a Gaussian noise (*µ*=0,*σ*=1). However, vector autoregression (VAR) is used to control the underlying dynamics of each sample. VAR explains the evolution of a variable over time with the generalized equation: *x*_*t*_ = *c* + *A*_1_*x*_*t*−1_ + *A*_2_*x*_*t*−2_ + … + *A*_*p*_*x*_(_*y − p*) + *e*_*t*_. For all samples, the VAR is computed using a positive semi-definite matrix, **A**. Then, 15 successive steps are randomly chosen to be perturbed with new dynamic information. Or, two more positive semi-definite matrices, B and C, are created and VAR is again used to compute 15 new steps using Gaussian noise and either matrix B or C, depending on the class label of the sample. These new steps, 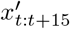 are added to the sample at a randomly selected interval (*x*_*t*:(*t*+15)_), with an interpolation variable, *α* resulting in the equation: 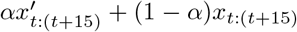. Examples of this data are seen in figure 5. This VAR dataset is again built specifically to show, without doubt, the efficacy of the methodology. However, in this case, as it is specifically engineered to highlight dynamical information, we argue that it is somewhat representative of fMRI data, where dynamical patterns are prevalent and highly influential.

**Figure 5:**
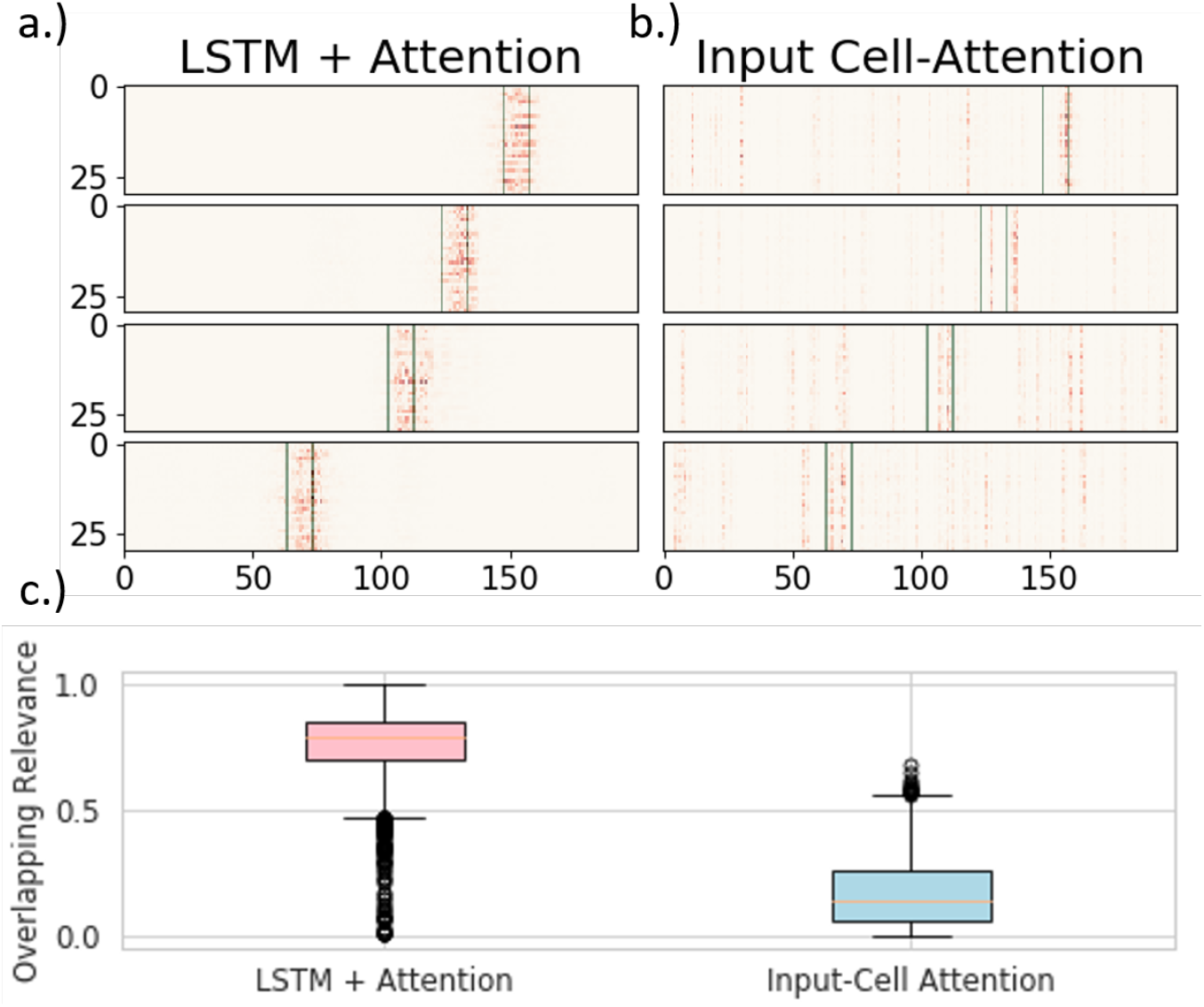
The results from the analysis of the VAR dataset. Several examples of resulting maps from both LSTM+attention (a) and input-cell attention (b) where the green rectangle is a mask representing the truly relevant information (i.e. where the auto-regressive signal changes). Boxplots (c) of the overlapping-values metric over all 3,000 holdout samples for both models. The overlap is defined as the percentage of the total sum of the maps that is within the relevant area over the total sum of the map. The green blocks in (a) and (b) highlight these relevant regions. Each baseline image has a certain underlying transition matrix, as computed by VAR, and each sample is interpolated with one of two different transition matrices, depending on the class label (the label along the y axis) within the highlighted relevant area. Notice the difficulty in which it can be to visually determine where the relevant information occurs. This gives the dataset a complexity that is not found in other, simpler datasets, including our boxes experiment.

### 2.6 Saliency Quality Metrics

Since the relevant information of the synthetic datasets is easily quantifiable, we can use basic similarity scores between the saliency maps and proper representations of the input to understand the quality of the maps. We compare our map quality on holdout samples to those of a input-cell attention model. Firstly, as the input data is noisy, we need a reasonable representation of each sample. For both experiments, we represent each sample as a binary matrix in which only the elements within the perturbed regions are ones, and all other elements are zero. In the first experiment, the randomly selected window of 10×15 elements are ones, and in the second experiment, the 15×30 window perturbed with added dynamics is our non-null region. Additionally, for the saliency maps from both our method and input-cell attention, we pass each sample through an absolute function [25]. To conduct a fair comparison with [7], we use both of the similarity metrics therein: Euclidean distance and weighted Jaccard similarity. We also evaluate the sum of all salient values within the window over the sum of the entire map.

To ensure an unbiased sampling of the timecourses with our model, we separate the data into 27000 training samples and 3000 test samples. We train 300 models on the non-holdout set and generate the maps for every sample, then select the saliency maps using the selection criteria described in our approach section and generate the saliency maps for the holdout set as well. We chose 300 due to computational restrictions, as each model can take some time to train. These maps are then fed through either a rectified linear unit (ReLU) function or absolute function (depending on the experiment) to avoid relying on both positive and negative derivatives to find the relevant information. Recent research has shown that removing negative values entirely from saliency can be beneficial [26].

### 2.7 Salient Networks

With the selected saliency maps, we sum along the temporal axis for each subject, resulting in a vector of size 53 for each subject. We then compute the group-wise sex differences for each component using Cohen’s D. We select Cohen’s D because, due to the large sample size of our data, we prefer an effect size measure that is agnostic to sample size. This analysis highlights which networks (and which brain regions) are significantly more important for the correct classification of men vs. women.

### 2.8 Co-Saliency

In order to better understand the sex differences within the ICA TCs, we computed the pairwise correlation of the processed saliency maps, which we call “co-saliency”, using Pearson correlation. These correlation matrices describe the relationships between the relevancy of time-varying values of the ICA components. Notably, they capture similar relationships to those found in fMRI connectivity.

## 3 Results

### 3.1 Synthetic Verification

Results from synthetic data show that our method is very effective at finding the truly relevant information. The weighted Jaccard and Euclidean distance showed vast improvement for our method over a current state-of-the-art method, input-cell attention for both the stationary dataset (in figure 4) and the more dynamic, VAR-induced dataset (in figure 5). The VAR dataset is especially significant as it shows that our method can properly capture non-stationary information. It is also important to note that both our method and the input-cell attention model got 99% accuracy on holdout data from the stationary dataset. However, our model achieved much higher accuracy (92%) on holdout data from the VAR dataset than input-cell attention (81%). Two tables of all metrics for both the boxes and VAR datasets are in table 1.

**Table 1:** Table comparing the LSTM+attention with the input-cell attention methodology on the boxes and VAR datasets. Within each cell is the average and standard deviation of the metric over 3,000 test samples, and the p-value comparing the two methodologies using a 2-sample t-test.

|  | Boxes Dataset |  |  | VAR Dataset |  |  |
| --- | --- | --- | --- | --- | --- | --- |
|  | Euclidean Distance | Overlapping Values | Weighted Jaccard | Euclidean Distance | Overlapping Values | Weighted Jaccard |
| LSTM + Attention | $\mu=1.24$,<br>$\sigma=.23$,<br>$p<.0001$ | $\mu=.33$,<br>$\sigma=.07$,<br>$p<.0001$ | $\mu=.22$,<br>$\sigma=.04$,<br>$p<.0001$ | $\mu=4.57$,<br>$\sigma=1.16$,<br>$p<.0001$ | $\mu=.56$,<br>$\sigma=.31$,<br>$p<.0001$ | $\mu=.10$,<br>$\sigma=.05$,<br>$p<.0001$ |
| Input-Cell Attention | $\mu=2.35$,<br>$\sigma=.45$,<br>$p<.0001$ | $\mu=.15$,<br>$\sigma=.05$,<br>$p<.0001$ | $\mu=.07$,<br>$\sigma=.02$,<br>$p<.0001$ | $\mu=5.34$,<br>$\sigma=1.42$,<br>$p<.0001$ | $\mu=.17$,<br>$\sigma=.14$,<br>$p<.0001$ | $\mu=.05$,<br>$\sigma=.03$,<br>$p<.0001$ |

### 3.2 Sex-Relevant functional Activity

To verify that our model is learning discriminatory patterns, we use stratified 10-fold cross-validation across the entire dataset, and find that the average validation accuracy over all folds and all models was 91.3%. As we used 10-fold cross-validation, we accumulated the predictions for each subject when they were used as a test sample for each model. In the end, we had 2,464,800 predictions (300 models * 8216 subjects). 90.5% of women were correctly predicted, and 91.8% of men were correctly predicted. A confusion matrix is found in table 2. Our model’s performance can be compared to other fMRI studies, where accuracy can range from 85% accuracy [27] to 94% [28], showing our results fit well with the current spectrum. It should also be noted that prediction with structural MRI can lead to much higher accuracy, closer to 100% [1].

**Table 2:** The confusion matrix over the average of all models and folds (3000 in total) for the UKB classification accuracies using our modality. The values are normalized to be percentages.

|  | *Predict Female* | *Predict Male* |
| --- | --- | --- |
| *True Female* | .905 | .095 |
| *True Male* | .082 | .918 |

Figure 6 shows the components with the highest Cohen’s D effect size between groups based on the per-component relevancy averaged over time, where red networks are more salient for women, and blue are more salient for men. We find that the AUD domain is particularly relevant for women [29]. The SC and VIS domains also appear to be highly relevant for male classification, with the highest biological sex differences being within the VIS domain. Finally, the SM domain is highly relevant, with different networks signaling for the two sexes.

**Figure 6:**
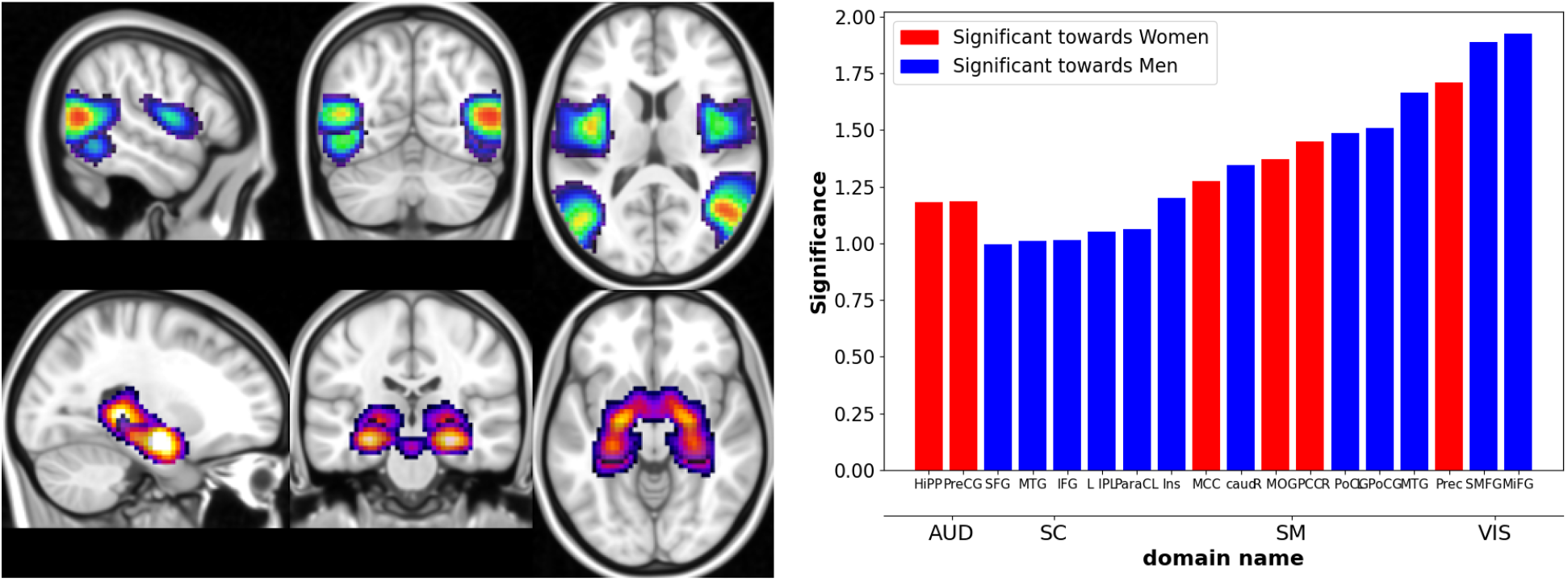
Two ICA components, the middle temporal gyrus (top), which is most significant for men, and the hippocampus, which is most significant for women (middle). Bar graph (right) showing the effect size of the most significantly different components between men and women. Blue components are directed towards men, and components in red are directed towards women.

### 3.3 Co-Saliency Analysis Differences

From the co-saliency maps we find that the differences are almost “orthogonal”, showing that the saliency method starkly separates the two sexesby focusing on network-specific timepoints that are intercorrelated in functionally structured ways. The co-saliencies indicate that temporal patterning of network-specific saliencies in women is significantly more strongly correlated than they are in men, as can be seen in the connectivity matrices in figure 5. This strong co-saliency for women suggests that the model is identifying timepoints in which networks are more tightly aligned when correctly classifying women, with networks in less tight temporal alignment at salient timepoints for the correct classification of men. Figure 7 highlights the co-saliency differences in comparison to the differences among static connectivity. Primarily what we see is that both co-saliency and sFNC show sex differences in the VIS, CC, and DMN domains.

**Figure 7:**
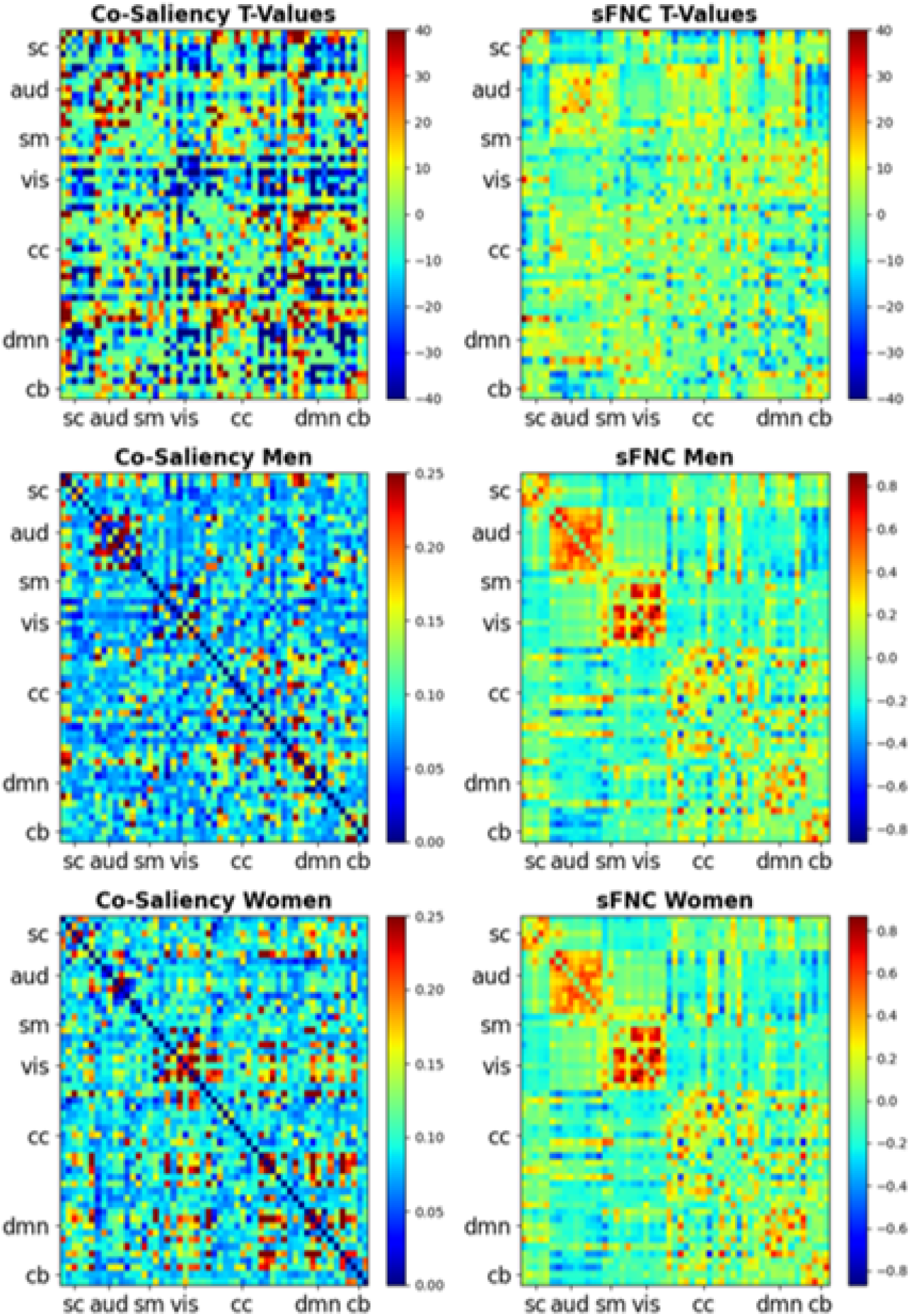
Heatmaps for the co-saliency (left) T-values masked by the FDR corrected significance, where *α* = .01 (top), the co-saliency for men (middle) and co-saliency for women (bottom). The corresponding sFNCs are on the right side.

We capture this modularity with greedy graph modularity estimations. In figure 8, we can see the estimated communities of the two sexes with the Clauset-Newman-Moore greedy algorithm [30]. From these communities, we computed the overall modularity of each sex [31], giving us a final modularity score of .37 for women, and .31 for men. The modularity for the sFNCs was .985 for women and 1.081 for men. The sFNC and co-saliency matrices organized by community are also found in figure 8.

**Figure 8:**
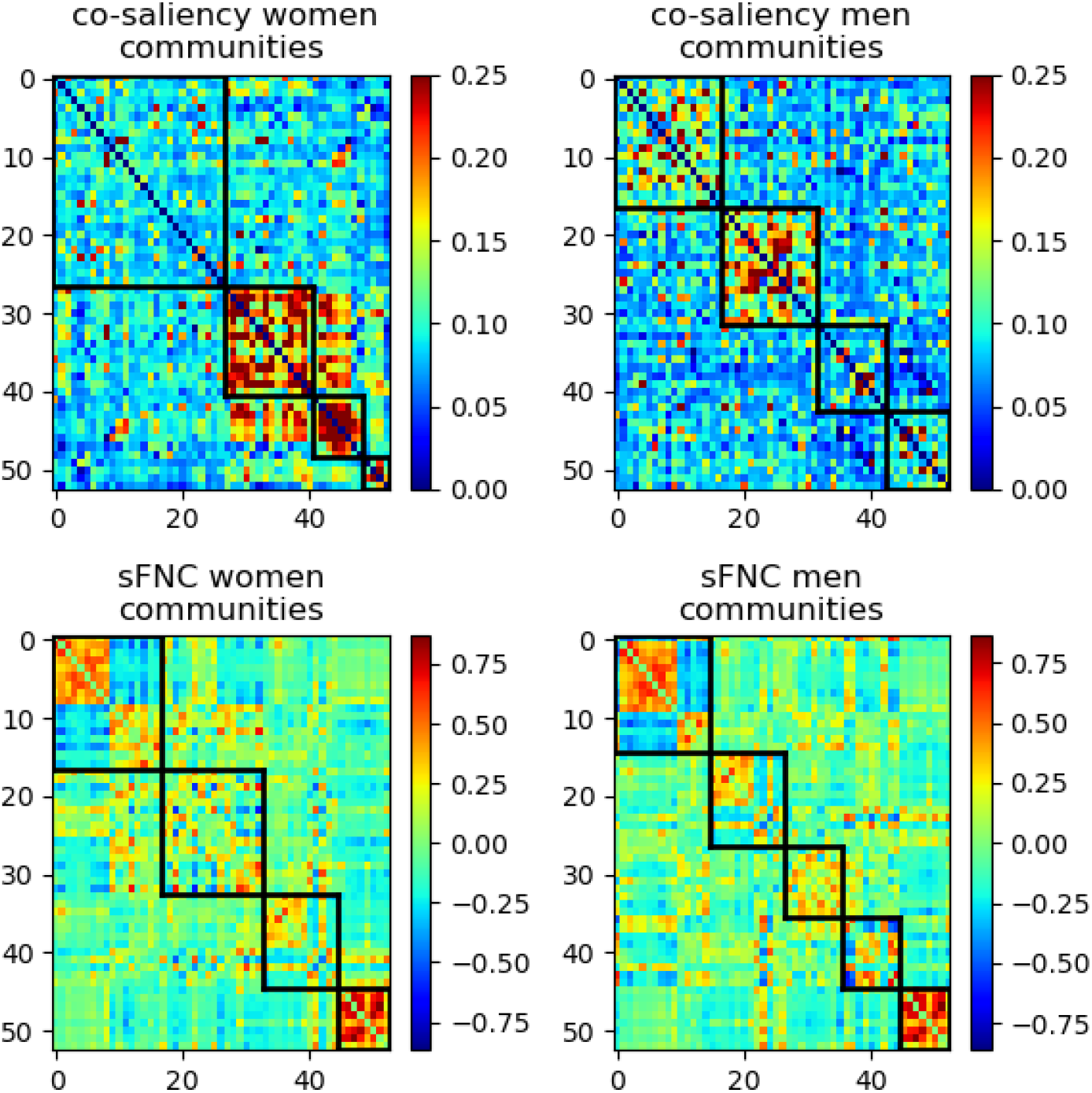
The co-saliency (top) and sFNC (bottom) matrices organized by communities (black boxes) for women (left) and men (right).

## 4 Discussion

After validating our methodology on synthetic data [22], we extract classification-relevant sex differences within the ICA timecourses. Overall, we see that the saliency maps and post-hoc analyses capture several patterns that are found within the data. The ICN differences from the saliency maps show that there are several domains that are key for sex classification, with many of these differences being backed by previous literature. The similarities between our results and past research is important, as these results are values for the ICNs averaged over time, which makes them more global, and presumptively, more representative of global, high-level differences.

Figure 6 highlights the ICN and domain differences. The high effect size of the difference between men and women in the AUD domain shows that this domain is particularly relevant for the model’s classification of women. Our result is supported by previous research [32], which shows major differences within the AUD domain between the two sexes. The SM domain is also highly relevant for women, which is also supported by previous research [33]. These findings are quite interesting, as they are the only domains that are primarily more relevant for women over men. A striking aspect of our findings is that the visual domain seems to be very relevant for how the model classifies and understands biological sex. We also see that 12 networks are significantly relevant for male classification, whereas only 6 networks are significantly relevant for female classification. This is important because, as each map is normalized to be a probability map, it suggests that the relevant information for female classification is more concentrated to fewer networks than male classification.

In addition to VIS, the SC domain is primarily relevant for correct classification of males. Currently, there is a wealth of research connecting the SC to differences between men and women, from both MRI and functional MRI studies [34]. Specifically, previous research has found that sex differences in the gyrus [35], which supports our findings. These previous findings show differences in complexity, or the spatial frequency of the brain surface [36].

Our co-saliency analysis, elucidated in figure 7, provides a wealth of information, much of which is consistent with previous results. The areas of consistency offer indirect validation of the relatively new method presented here, which is still being developed and refined. One important finding is that the dynamics of the networks within the CC domain, in relation to networks both within and outside of the CC domain, figure strongly in the model’s ability to differentiate male and female subjects. The connections between CC networks and the AUD, DMN, and SC domains significantly inform the model’s decisions. Many networks within the CC domain also play a major role in model classification, and we see a split, where some networks are highly relevant for male classification, and others are highly relevant for female classification. Specifically, connections within and between the inferior and middle-frontal gyrus and the hippocampal networks are primarily relevant for men. For women, the left inferior parietal lobule, the middle cingulate cortex, and the superior frontal gyrus all have especially salient connectivity patterns. These two domains, CC and VIS also show stark coherency differences, which can be visible in the co-saliency maps, and quantified in the community detection and graph-modularity computation seen in figure 8. The differences that are weighted towards men, however, appear de-modularized compared to the sFNCs. It’s also interesting to note that the most significant VIS co-saliency pairs are entirely within the domain and female-centric. These patterns also appear in previous connectivity work [37, 38]. The co-saliency maps seem to show these patterns, but with a great deal more contrast than in the raw data. This contrast enhancement highlights subtle differences that are only weakly evident in the raw data, differences that look negligible under a linear, univariate lens, but prove highly relevant to biological sex when employed in a multivariate nonlinear classification model. Our results also show several key findings that appear to have been missed by non-linear analysis. Specifically, we see substantial differences in how certain domains are organized. We find that VIS and DMN domains, as well part of the CC are prominently modular for women. From the community and general graph analyses, we also see an overall lack of modularity and even coherence for men. Figure 8 highlights our community/graph analysis results, which show the overall modularity differences. The findings we report are complex and do not admit straightforward interpretation framed by previous results. They suggest, however, that our understanding of biological sex has been limited by the ubiquitous use of linear univariate models, and that expanding the traditional model space could help better realize the scientific promise of noninvasive functional brain imaging.

The field of model interpretability is still young and rapidly evolving. While the focus of this paper is to introduce approaches to stabilize and quantify deep-learning interpretability methods for analyzing brain imaging data, new and more effective methods may emerge revealing different information. Recurrent models themselves, as we highlight in this paper, are architecturally challenging to interpret using gradient-based approaches. Although we suggest that our methods mitigate the gradient bias against distal timepoints and model instability across initializations, the computed saliencies remain sensitive to changes in hyperparameters and architecture, and overall they can still be underspecified.

Another impediment to model interpretability is the complexity of the data itself. Although we have many carefully crafted and effective methods for quantifying information from ICN timecourses, these timecourses are highly processed dimensional reductions of the original scan data. This adds another layer of complexity to understanding the brain functionality through the lens of saliency maps, increasing the possibility of erroneous interpretation of the findings. These flaws can be mitigated in due time. As more researchers focus on these topics, both specifically to neuroimaging and in general, we will find more robust and effectively interpretable maps. We also suggest that new methods to analyze the maps (such as co-saliency) will become more prominent, opening the door for a better understanding of intricate datasets, and especially neuroimaging.

